# Metatranscriptomic reconstruction reveals RNA viruses with the potential to shape carbon cycling in soil

**DOI:** 10.1101/597468

**Authors:** Evan P. Starr, Erin E. Nuccio, Jennifer Pett-Ridge, Jillian F. Banfield, Mary K. Firestone

**Affiliations:** Department of Plant and Microbial Biology, University of California Berkeley, CA, USA; Nuclear and Chemical Sciences Division, Lawrence Livermore National Laboratory, Livermore, CA, USA; Department of Earth and Planetary Science, University of California, Berkeley, CA, USA; Earth Sciences Division, Lawrence Berkeley National Laboratory, Berkeley, CA, USA; Department of Environmental Science, Policy, and Management, University of California, Berkeley, CA, USA; Chan Zuckerberg Biohub, San Francisco, CA, USA; Innovative Genomics Institute, Berkeley, CA, USA

**Keywords:** Virus, phage, soil, rhizosphere, metatranscriptome

## Abstract

Viruses impact nearly all organisms on Earth, with ripples of influence in agriculture, health and biogeochemical processes. However, very little is known about RNA viruses in an environmental context, and even less is known about their diversity and ecology in the most complex microbial system, soil. Here, we assembled 48 individual metatranscriptomes from four habitats within a soil sampled over a 22-day time series: rhizosphere alone, detritosphere alone, a combination of the two, and unamended soil (four time points and three biological replicates per time point). We resolved the RNA viral community, uncovering a high diversity of viral sequences. We also investigated possible host organisms by analyzing metatranscriptome marker gene content. Based on viral phylogeny, much of the diversity was *Narnaviridae* that parasitize fungi or *Leviviridae* that infect Proteobacteria. Both host and viral communities appear to be highly dynamic, and rapidly diverged depending on experimental conditions. The viral communities were structured based on the presence of litter, while putative hosts appeared to be impacted by both the presence of litter and roots. A clear time signature from *Leviviridae* and their hosts indicated that viruses were replicating. With this time-resolved analysis, we show that RNA viruses are diverse, abundant and active in soil. Their replication causes host cell death, mobilizing carbon in a process that represents a largely overlooked component of carbon cycling in soil.

## Introduction

We have much to learn about soil, the world’s most diverse and enigmatic microbial habitat. With the advent of meta-omics techniques, the diversity, ecology, and impact of bacteria and archaea in soil is being rapidly revealed (1). Recent work on double stranded DNA bacteriophage (phage) in soil has begun to explore their diversity and host interactions (2, 3). However, other members of soil communities, such as RNA-based phage, RNA viruses and eukaryotes have not been as deeply explored using meta-omic methods. RNA viruses are the counterpart to DNA viruses, differing in the nature of their genetic material and less studied in environmental contexts because they are not captured in DNA sequencing studies. In some systems, the impact of RNA viruses on community and ecosystem processes has been proposed to rival or exceed the impact of DNA viruses (4). The majority of soil RNA viral work has been single-host focused, for agriculturally relevant crops (5) or crop pathogens (6). Environmental RNA virus studies have focused on less complex or more tractable systems such as marine environments and the human and animal gut (7–11). To our knowledge, no sequencing-based RNA viral community analyses have investigated soil.

Viruses are major players in biogeochemical cycling. However, much of what is known about viral impacts on elemental cycling comes from aquatic systems. Marine phages can lyse up to one-third of bacteria in ocean waters per day, releasing a huge amount of carbon (12–14). The released components include dissolved organic carbon (DOC) that is readily metabolized by heterotrophic bacteria, but largely inaccessible to eukaryotic grazers and higher trophic levels (15). This phenomenon is termed “the viral shunt” (15, 16), and is thought to sustain up to 55% of heterotrophic bacterial production in marine systems (17). However, some organic particles released through viral lysis aggregate and sink to the deep ocean where they are sequestered from the atmosphere (18). Most studies investigating viral impacts on carbon cycling have focused on DNA phages, even though up to half of all viruses in the ocean may be RNA viruses (4). By analogy, lysis of organisms by RNA viruses may represent a large contribution to carbon flow in soils. The process of cell lysis and carbon liberation likely stimulates mineralization of biomass and contributes to carbon stabilization when compounds are sorbed to mineral surfaces or occluded within aggregates (19, 20).

Here, we used assembled metatranscriptomic data from a California annual grassland soil to reconstruct RNA viral genome sequences and constrain their possible hosts. We searched the assembled sequences for the RNA-dependent RNA polymerase (RdRp), a conserved gene in all RNA viruses that lack a DNA stage (generally referred to as Baltimore type III-V RNA viruses). Although recent work tracing the deep evolutionary history of RNA viruses has merged viral families into supergroups (21–23), our focus was not on resolving deep phylogenetic viral branches. Rather, our objectives were to investigate fine scale diversity of RNA viruses, their putative hosts, and possible functional importance in soil. Thus, we relied heavily on Shi et al. (2016), the most comprehensive metatranscriptomic study of RNA virus diversity so far, for taxonomic classification and to delineate viruses from previously undocumented clades.

The diversity of soil eukaryotes that may serve as hosts for RNA viruses remains largely understudied. Generally, RNA viruses are known to infect fungi, plants, animals and the many clades of single-celled eukaryotes, in addition to some Proteobacteria. Soil eukaryotic studies have heavily relied on primer based sequencing or visual classification, methods that can impart biases and miss novel organisms. Using the genomic information contained in our assembled metatranscriptomes, we identified many clades of eukaryotes without reliance on primers or microscopy. We tracked both viral and eukaryotic communities in key soil habitats: the rhizosphere, detritosphere (the area of influence around dead particulate organic matter), a combination of the two, and unamended soil (bulk soil) over time. The relatively short sampling time scale (3, 6, 12, and 22 days) allowed us to investigate viral and host community dynamics. Our research places direct constraints on the timescales for virus dynamics and provides the first genomic view of RNA viruses in soil.

## Results

### Experimental design

Wild oat (*Avena fatua*), an annual grass common in Mediterranean climates, was grown in microcosms with a sidecar, allowing us to track root age and sample from the rhizosphere (24). The experimental setup involved microcosms, half of which included soil mixed with dried ground *A. fatua* root litter and the other half contained soil without litter amendment. All microcosms contained bulk soil bags, which excluded roots. Once *A. fatua* was mature, roots were allowed into the sidecar and the growth of individual roots was tracked. We destructively harvested rhizosphere soil (and paired bulk soil) that had been in contact with the root for 3, 6, 12, and 22 days. In total, we sampled paired rhizosphere and bulk samples from four time points with two treatments (with and without litter), with three biological replicates, for a total of 48 samples for metatranscriptome sequencing. We sequenced a total of 408 Gbp with an average of 8.7 Gbp per sample.

### Eukaryotic RNA viruses

We used profile hidden Markov models (HMM) to search our assembled metatranscriptomes and found a total of 3,884 unique viral RdRp sequences (dereplicated at 99% amino acid sequence identity; AAI). This includes 1,350 RNA bacteriophage (phage), viruses that infect bacteria, and 2,534 viruses which infect eukaryotes.

Our eukaryotic viruses group into 15 major clades that span the majority of known viral diversity (**Fig. 1**). Many were included into the supergroup-like clades defined by Shi et al. (2016). For the remainder, we constructed phylogenetic trees to define additional viral families.

**Figure 1.**
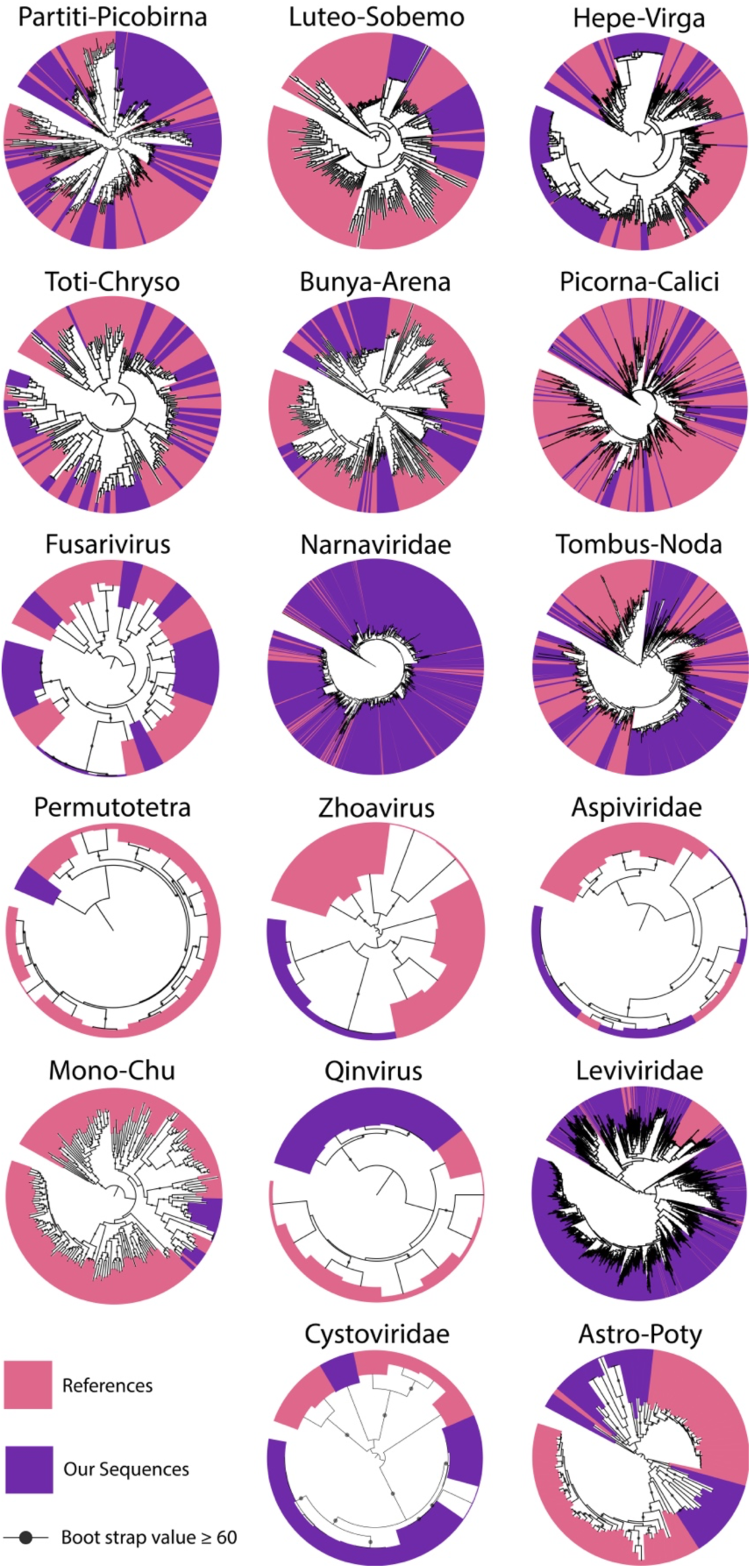
Phylogenetic trees representing clades of viruses based on RdRp. Within each tree the RdRp sequences we identified are colored purple and previously described sequences are in pink. Trees are all midpoint rooted. More detailed trees are shown in Supplementary Data.

Overall, in trees that include both existing and newly generated sequences, we noted a strong grouping of our RNA viral sequences into ‘fans of diversity’, much like the seed head a dandelion. Many of these fans included a single reference sequence previously used to propose a new viral family. For example, in the Hepe-Virga, sequences grouped with the *Agaricus bisporus virus 16* (proposed family *Ambsetviridae* (25), in the Astro-Poty clade, sequences group with *Bufivirus UC1* from wastewater (22) and in the Partiti-Picobirna, sequences cluster with *Purpureocillium lilacinum nonsegmented virus 1* (26). We substantially expanded the *Barnaviridae* (Lueto-Sobemo superfamily) and *Mymonaviridae* (Tombus-noda superfamily) families (27, 28), which replicate in fungi, and the newly proposed *Zhaovirus* and *Qinvirus* families, which are found in invertebrates (23).

We predict that fungi are the most common hosts for many of our newly reported RNA viruses (mycoviruses; **sup fig. 1**). We are most confident when they fall into the *Barnaviridae, Megabirnaviridae, Quadriviridae*, and mitoviruses, groups currently thought to only infect fungi (29, 30). The most frequently encountered virus in our dataset, accounting for over 50% of the eukaryotic viral strains identified, came from the mitovirus genus in the *Narnaviridae* family. Mitoviruses are linear single stranded RNA viruses that replicate within fungal mitochondria and spread vertically through spores and horizontally through hyphal fusion (31, 32). We suspect that most of the mitoviruses we detect are infecting fungi because, although some mitoviral sequences have been found integrated into the genomes of plants, they are frequently truncated and not transcribed (33, 34). Mitoviruses were also the most abundant viral clade in every sample (**sup fig. 2**), and one specific mitovirus was ∼30X more abundant that the most abundant fungus in the same sample, based on coverage.

**Figure 2.**
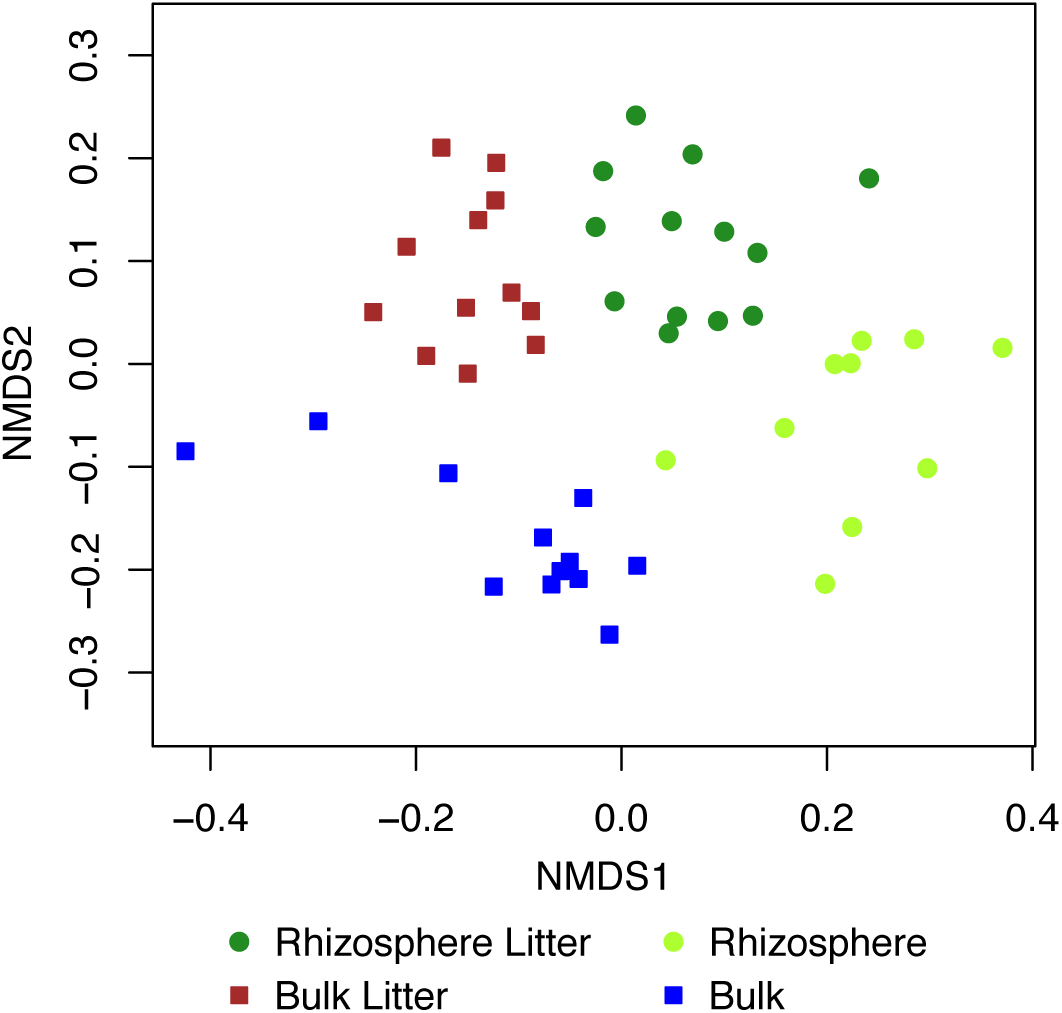
Nonmetric multidimensional scaling ordination of eukaryotic communities based on coverage of the Cox1 gene.

Until recently, the *Narnaviridae* group, which includes mitoviruses, were thought to only encode an RdRp, but a recent discovery suggests some narnaviruses (only distantly related to mitoviruses) encode additional proteins, including capsids and helicases (23). The majority of mitoviruses we identified contained only a single RdRp gene. However, we predicted several additional proteins on some mitoviral genomes (**sup fig. 3**). These putative genes are small (average 79 amino acids) and novel, and functions could not be predicted for them. Sometimes the additional gene(s) are transcribed in the same direction as RdRp and in other cases they are transcribed in the opposite direction (**sup fig. 3**).

**Figure 3.**
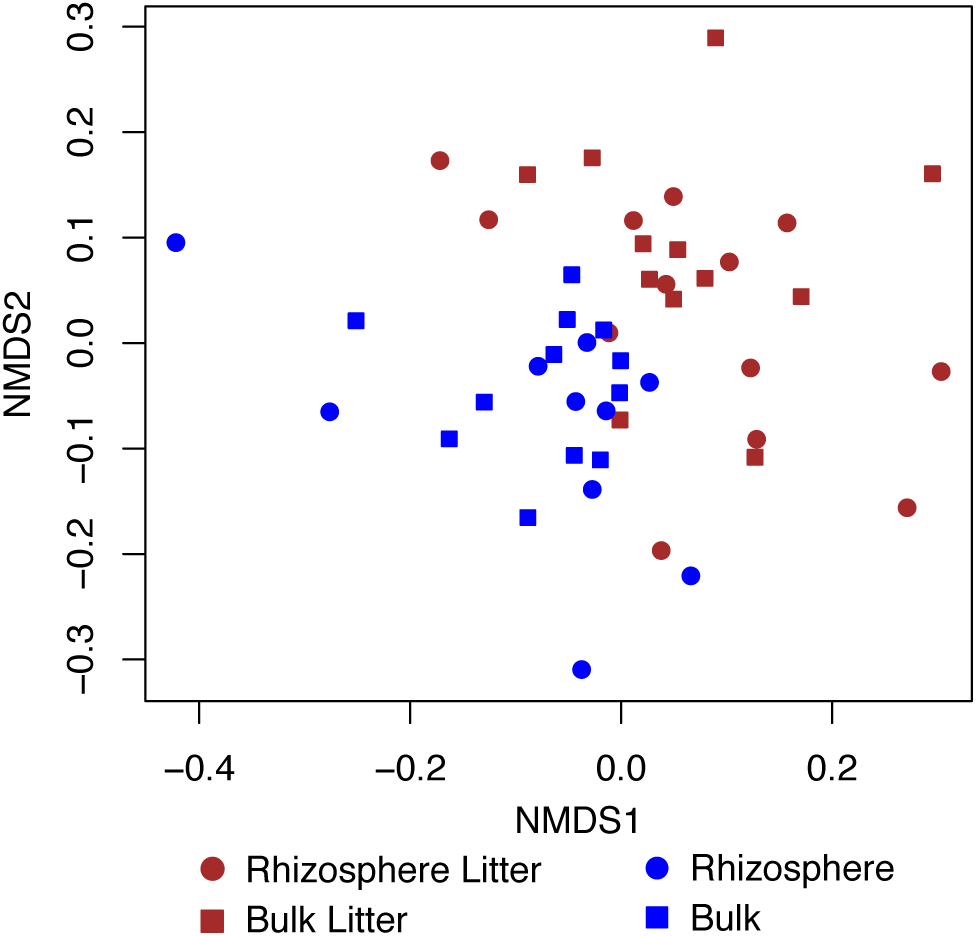
Nonmetric multidimensional scaling ordination of eukaryotic viral communities based on coverage of the viral scaffolds.

We reconstructed many sequences for viruses that likely infect eukaryotes other than fungi. In the Picorna–Calici, the hosts are likely vertebrates, insects, algae, and plants based on viral phylogenetic placements (35). In the Tombus-Noda tree, many of the RdRps group with umbraviruses, well recognized plant viruses. Other RdRp sequences group with sequences from complex environmental samples, but the hosts are unknown.

RNA viruses from some previously defined superfamilies were conspicuously absent. We did not identify any soil RNA viruses belonging in the *Nidovirales*-like, *Reoviridae*-like and *Orthomyxoviridae*-like superfamilies. The absence of *Reoviridae* is interesting, as these RNA viruses can infect fungi (36). We also did not confidently identify any *Ortervirales*, retroviruses that replicate with a DNA intermediate. We identified many reverse transcriptase proteins, but none of the corresponding scaffolds encoded capsid proteins, so the sequences may be retrotransposons or fragments of retroviruses. We classified two new clades within the Bunya-Arena viral super-family as novel, given that they exhibit sequence divergence comparable to that which separates known RNA viral families (see supplementary information). In both cases, the scaffolds only encode the polymerase with a Bunyavirus RNA dependent RNA polymerase domain. This genome structure is shared by *Hubei myriapoda virus 6* (23) and *Ixodes scapularis associated virus 3* (37).

### Eukaryotic hosts

As viruses cannot replicate without their host, changes in virus abundance levels implicate introduction of new vectors or hosts, shifts in host abundance levels, or changes in host susceptibility to infection. To better understand the diversity and ecology of RNA viral hosts and carriers in soil, we used the mitochondrially encoded cytochrome C oxidase subunit 1 (Cox1) gene as a marker to define the eukaryotic populations present (38–43). However, mitochondrial transcription is determined by metabolic activity of a cell (44), and can be impacted by the switch to a symbiotic state (e.g. in arbuscular mycorrhizal fungi (45)) or nutrient sensing and hormonal signals (in animals (46)). Thus, the number of reads mapped to the Cox1 genes is determined by the number of organisms present and organism activity, so it is an overall measure of the importance of each organism in the community.

We identified 726 eukaryotic Cox1 sequences and clustered them at 98% AAI to approximate a species level view. The eukaryotes we discovered span the diversity of known soil organisms. Not surprisingly, sequences from *A. fatua* were the most abundant Cox1 transcripts in many samples. In some samples, the most abundant Cox1 were from an *Enchytraeidae* related worm or an *Amoebozoa sp.* Other samples were dominated by a mixture of fungi, Amoebozoa, Viridiplantae, and unknown eukaryotes.

Amoebozoa were the most diverse eukaryotic clade in this dataset, >25% of the identified eukaryotes. Next most diverse were species of fungi and unknown eukaryotes. The many unknown eukaryotes reflects the lack of environmental Cox1 sequences from micro-and meso-eukaryotes in public databases. For this reason, we reconstructed 18S rRNA gene sequences for classification, as they are better represented than Cox1 in public databases. Although transcripts had been depleted in bacterial and plant ribosomal RNA, a sufficient amount of 18S rRNA sequences persisted that we could identify 521 distinct species (after clustering at 98% nucleic acid identity to approximate species groups; **Table 1**). The results revealed the presence of Centroheliozoa, Malawimonadidae and Jakobida not identified on the Cox1 analyses and many more species of Heterolobosea, Euglenozoa and Rhizaria.

**Table 1.** Number of eukaryotes identified based on marker genes. *Note that these results may be skewed by rRNA depletion.

| Clade | Cox1 | 18S rRNA* |
| --- | --- | --- |
| Amoebozoa | 193 | 295 |
| Fungi | 174 | 33 |
| Unknown eukaryotes | 108 | 14 |
| Metazoa | 85 | 28 |
| Stramenopiles | 55 | 17 |
| Alveolata | 54 | 14 |
| Heterolobosea | 11 | 41 |
| Nucleariidae | 11 | 0 |
| Cryptophyta | 9 | 1 |
| Euglenozoa | 7 | 28 |
| Viridiplantae | 6 | 2 |
| Choanoflagellida | 5 | 8 |
| Rhizaria | 3 | 21 |
| Apusozoa | 2 | 1 |
| Alveidia | 2 | 0 |
| Ichthyosporea | 1 | 0 |
| Centroheliozoa | 0 | 11 |
| Malawimonadidae | 0 | 6 |
| Jakobida | 0 | 1 |

### Eukaryotic virus and host ecology

Both litter and rhizosphere habitats shape the eukaryotic community structure, as measured by the Cox1 gene transcript abundance (**Fig. 2**). Samples split based on presence of the root and litter into four clusters: no-litter, root and litter, bulk soil with no-litter, and soil with litter. Separation of the communities was evident by the time of the first sampling (3 days) and persisted to the end of the 22-day long experiment. However, when the *A. fatua* Cox1 sequences were removed from the dataset (**sup fig. 4a**) there was much less community separation between rhizosphere and bulk soil (PERMANOVA R^2^ 0.17, P< 9.999e-05 to R^2^ 0.03, P< 0.005) and a time signature was detected (**sup fig. 4b**).

**Figure 4.**
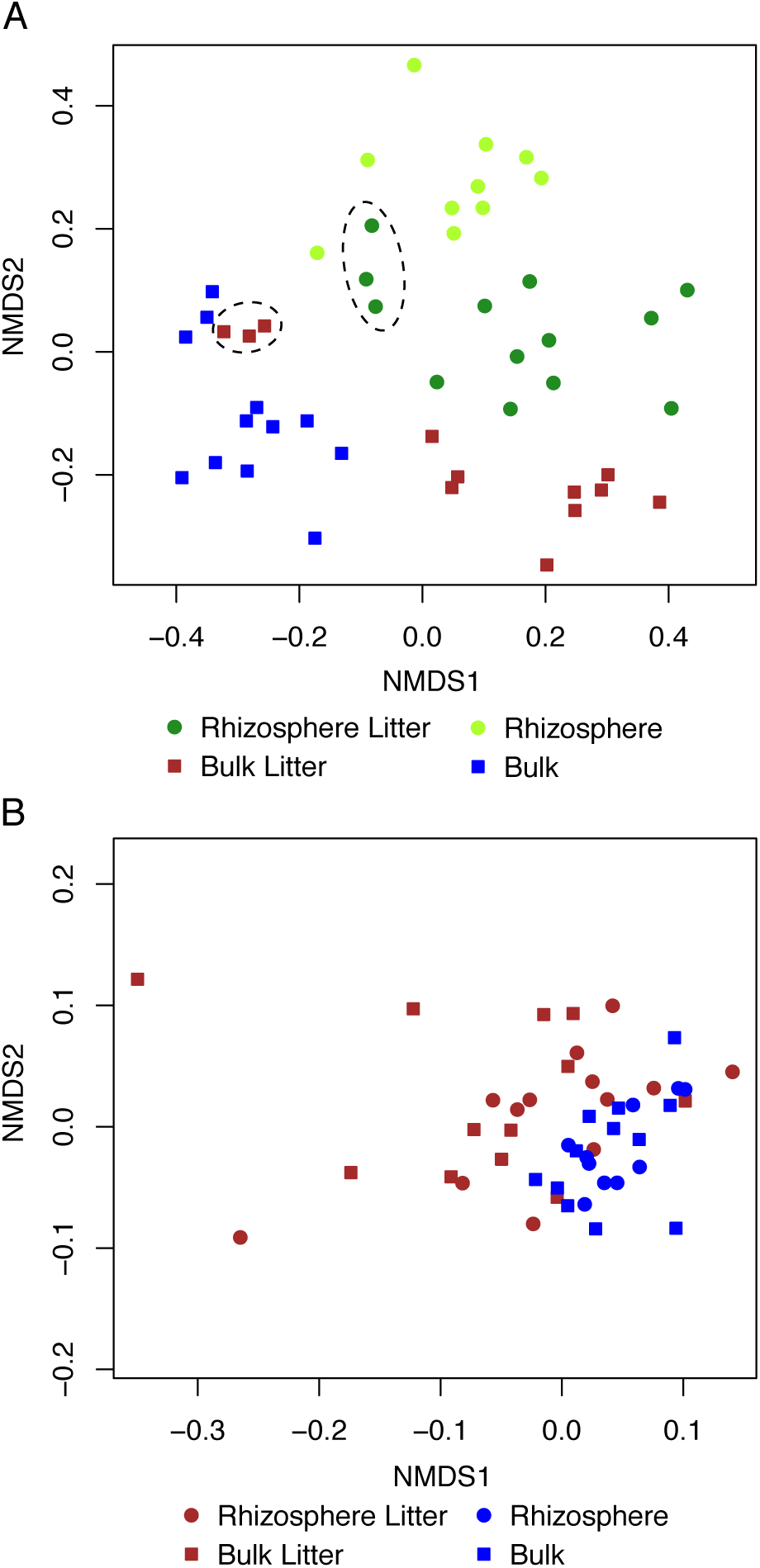
Nonmetric multidimensional scaling ordination of the Proteobacterial community (A) and the *Leviviridae* community (B). Dashed ellipses in (A) are litter amended samples (bulk with litter and rhizosphere with litter) at day 22.

We compared the abundances of eukaryotic species in unamended bulk soil to their abundances in soils that included the rhizosphere and litter, soil with litter, and rhizosphere without litter to identify species statistically enriched in each case (**sup fig. 5**). The results showed that the presence of litter and rhizosphere enriched for many Amoebozoa and fungi.

However, enrichment patterns indicate that litter had a greater selective force on more individual species than the presence of the root.

Root litter had a significant effect on the eukaryotic RNA viral community (PERMANOVA P< 9.999e-05); the presence of growing roots had no detectable impact (PERMANOVA P< 0.07; **Fig. 3**). The distinction between the litter vs. no litter samples was evident within 3 days, implying that the viral and eukaryotic communities changed at similar rates. Differences persisted to the end of the 22-day long experiment. The same patterns were replicated when the *Narnaviridae* were removed from the dataset, indicating that the numerous, unusual host-bound viruses, were not causing the previously identified trends (**sup fig. 6**).

We did not measure an effect of growing roots on viral abundances. However, a *Luteovirus* (99% AAI to Barley yellow dwarf virus PAV, which can infect *Avena* (47)) was the most abundant virus detected in one rhizosphere-litter sample. We identified more eukaryotic RNA viruses than eukaryotes, however twice as many eukaryotes showed a statistically significant enrichment under one or more condition (**sup fig. 7**). This may reflect the great heterogeneity in viral abundance patterns within replicates, as ~ 2% of the eukaryotic RNA viruses were found in only one sample compared to 0.1% of eukaryotes. The viral strains most frequently enriched relative to bulk soil were from the fungal mitochondria-associated *Narnaviridae* family, whose abundance is likely closely linked to the abundance and activity of fungi that responded strongly to the treatments.

### RNA phage, potential hosts and ecology

Two families of RNA viruses are known to infect bacteria, the *Cystoviridae* which infect *Pseudomonas sp.* (48, 49) and the *Leviviridae*, which infect Gammaproteobacteria and Alphaproteobacteria (50–52). We used a marker gene approach to find the RdRps of these RNA phage in the assembled transcripts. After dereplication, we identified 12 *Cystoviridae* sequences and 1,338 unique *Leviviridae* RdRp sequences. To put this into context, there are currently just over 200 *Leviviridae* RdRp sequences in public databases. Some of our sequences group with allolevivirus and levivirus, well-studied genera, but importantly, other *Leviviridae* sequences resolved new clades (**Fig. 1**).

The specific *Leviviridae* enriched in bulk soil are phylogenetically novel (**sup fig. 8**). Similarly, the most abundant *Leviviridae* in 46 of 48 samples places in a region of the tree lacking reference sequences. The 4,668 bp genome of this abundant RNA phage encodes four, nonoverlapping genes and has unusual genome architecture (**sup fig. 9 top**). Like *Enterobacteria phage M*, it may also encode a +1 frameshift lysis protein inside of the RdRp gene (53). The predicted lysis protein contains a gram-negative signal peptide and a transmembrane domain and may form pores in the plasma membrane, dissipating proton motive force and inducing cell autolysis (54). A related Leviviridae appears to encode 5 genes and possibly two frameshift lysis proteins (**sup fig. 6 bottom**). However, biochemical work is needed to resolve the function of these predicted proteins.

We used ribosomal protein S3 (rpS3) phylogeny to identify members of the bacterial and archaeal communities that may serve as hosts for *Leviviridae*. After dereplication at 99% AAI, we identified 717 species, 355 of which are within the Proteobacteria. Based on prior research, these are likely hosts for the *Leviviridae*, although other hosts cannot be ruled out. These proteobacterial abundances separate with rhizosphere, bulk, rhizosphere with litter, and bulk with litter treatments, much like the eukaryotic communities (PERMANOVA P< 9.999e-05; **Fig. 4a**), and this was evident after three days. By the final sampling point (22 days) the bulk with litter samples clustered with the bulk samples and the rhizosphere with litter samples clustered with the rhizosphere, indicating the disappearance of a discreet litter community (**Fig. 4a**). The *Leviviridae* communities also diverged within three days and showed a time signature (PERMANOVA P< 0.02; **sup fig. 10**). The communities separated based on the presence or absence of litter (PERMANOVA P< 9.999e-05), with root influence being undetectable (PERMANOVA P< 0.09; **Fig. 4b**).

## Discussion

To our knowledge, no prior study has genomically investigated the ecology and diversity of RNA viruses in soils. Using assembled metatranscriptomes, we uncovered a vast diversity of RNA viruses (>3,000 sequences) from a single grassland soil and discovered several possible new families. Some viruses grouped phylogenetically with viral families previously proposed based on only a single member, indicating soils may hold much of the diversity of these understudied groups. The viruses we reconstructed came from nearly all known groups of RNA viruses. We identified hundreds of eukaryotes and bacteria that may represent RNA viral hosts or vectors that can pass infections among plants and other organisms. This transfer of viral agents can have devastating and economically relevant effects (55–57).

Many of the viruses identified group phylogenetically with fungal viruses (mycoviruses), and fungi appear to be one of the main viral hosts in the studied soil. The most diverse mycoviruses are mitoviruses (*Narnaviridae*) which, despite their simplicity, can have significant impacts on fungal fitness and physiology (32, 58, 59).

The eukaryotic RNA viral and eukaryotic communities were strongly impacted by the soil experimental variables, especially the presence of root litter. By the first sampling timepoint, the eukaryotic and eukaryote-associated RNA viral communities were different in the presence and absence of root litter. Saprotrophic fungi likely responded favorably to the dead litter biomass. In contrast, dissolved organic compounds exuded by roots may be less accessible to non-fungal soil eukaryotes. Accordingly, roots had little impact on the eukaryotic community composition. Addition of dead plant biomass promoted the activity of detritivores, which likely immigrated into the litter from surrounding soil, increased transcription levels, grew, and/or germinated from spores. However, shifts in the eukaryotic RNA viral community are likely due to proliferation following increased activity of their hosts, although they may have also been transported by vectors.

RNA phage, which replicate in bacteria, are relatively little studied across all environments. Here, we provide the first glimpse into their ecology and diversity in soil. Given that over half of all identified RdRp sequences were *Leviviridae*, soils generally may prove to be hotbeds for *Leviviridae* diversity. As *Leviviridae* do not have a known lysogenic life stage, most of their population changes likely reflects replication in their hosts. Within the first 3 days of the experiment, the Leviviridae communities and their hosts were distinctly different indicating that these RNA viruses infected and replicated within days. Infections over this timescale likely have drastic effects on their host communities, and thus soil ecology.

In general, the magnitude of viral impacts on the soil carbon cycle is underexplored. Little research has been done on phage-induced bacterial lysis in soil and even less on viral-induced death of fungi and other eukaryotes, which can contain an equal or greater biomass compared to bacteria in some soils (60). The high diversity and abundance of identified RNA viruses, combined with their dynamic population changes, indicates that there was substantial viral replication. We hypothesize that proliferation of these lytic RNA phage and RNA viruses will have substantial impacts on the form, abundance and distribution of carbon compounds in soil. Lysis of host bacterial cells and viral-induced cell death of eukaryotes will release dissolved low molecular weight carbon compounds. These will likely be quickly consumed by nearby bacteria and much of the carbon returned to the atmosphere, mostly in the form of CO_2_. However, a portion of the cellular and viral debris and soluble carbon released may be stabilized and sequestered in the soil. For example, bacterial and eukaryotic lipids and polysaccharides could adhere to clay and other mineral surfaces or become occluded within soil aggregates.

RNA viruses and RNA phage were heterogeneously distributed across samples, including replicates. In contrast, the eukaryotes and bacteria appeared to have more even abundances across samples. In combination, these observations suggest that virus and phage abundances are not solely determined by the presence of their hosts, but factors such as sporadic blooms, probably in part due to variation in viral resistance levels in the host population, lead to patchy distribution patterns for viruses in soil. These patterns are analogous to those documented in marine virus blooms, though the exact cause of these events are still unknown (11, 61). Understanding the diversity and ecology of soil viruses may contribute to advances in biotechnology. Viral genomes have been mined for biopesticides and self-assembling nanomaterials (62, 63). Viruses have been proposed, and used, as biocontrol agents for culling invasive organisms including fire ants, rabbits and moths (64–66). Viruses are also being investigated as biocontrol agents for devastating plant pathogens such as, *Fusarium sp., Botrytis cinereal* and *Rosellinia necatrix* (6, 31, 67–71). Novel, environmentally derived viruses may be a source for new biotech tools and biocontrol agents.

## Conclusions

By reconstructing soil metatranscriptomes drawn from multiple soil habitats and informed by experiments we greatly increased the known diversity of RNA viruses. Phylogenetic analyses suggest that fungi are the most common hosts for RNA viruses in the studied grassland soil. The hosts for RNA phage remain to be definitively identified, but likely include Proteobacteria. Shifts in eukaryote, RNA phage and RNA viral abundances over a few day period reveal that entire soil communities can rapidly respond to altered resource availability. Our experiments indicate that the form of carbon inputs (root-derived low molecular weight C inputs versus macromolecular carbon compound in litter) may impact eukaryotic and bacterial abundance patterns, and that these in turn may be a major determinant of RNA viral and RNA phage dynamics.

## Declarations

## Supporting information

Supplementary Data 1. Tree files for each viral family.

Supplementary Data 2. Eukaryotic COX1 tree files.

Supplementary Data 3. 18S rRNA tree files.

Supplementary Data 4. Proteobacteria rpS3 tree files.

## Acknowledgements

This work was made possible by the support and expertise of the personnel at the Hopland Research and Extension Center (Hopland, CA, USA). Plants were grown at the UC Berkeley Oxford Tract Greenhouse Facility. Sampling efforts and technical expertise were provided by Shengjing Shi, Donald Herman and Katerina Estera-Molina. Methods and analysis input was provided by Ella Sieradzki and Alexander Probst.

## Contributions

EEN designed the experiment, with input from MKF. EEN setup the experiment, sampled, and prepared the RNA. EPS and EEN analyzed the data with support and guidance from JP. The manuscript was written by EPS and JFB and approved by all coauthors.

## Data availability

The sequences used in this study will be available on NCBI upon acceptance of the manuscript. Read files can be accessed through the JGI genome portal using GOLD study ID: Gs0110148.

### Ethics approval and consent to participate

Not applicable.

### Consent for publication

Not applicable.

## Methods

### Experimental design

Wild oat (*Avena fatua*) was grown in microcosms with a clear sidecar designed to allow access and visual tracking of the soil and rhizosphere (24, 72). Experimental soil was collected from Hopland Research and Extension Center (Hopland, CA. USA). The collection site has been described previously (73). Chambers were filled with soil and seedlings were planted in the microcosms and grown for six weeks before the start of the experiment. Six days before the start of the experiment the divider separating the main chamber and the sidecar was removed and the sidecar was packed with soil. The litter amended microcosms received 0.4g of dried *A. fatua* root litter. Bulk soil was placed in 18 μm mesh bags which allowed solutes to pass but not roots.

### Sample Collection

The ages of individual roots were tracked to collect rhizosphere soil which had been influenced by the root for 3, 6, 12 and 22 days. Three replicate microcosms were destructively harvested for paired rhizosphere and bulk soil. Rhizosphere soil was cut out from the rest of the soil along the edge of the root hair zone (<2 mm from the main root). Root sections and adhering soil were placed immediately in ice cold Lifeguard Soil Preservation Reagent (MoBio) in 2 ml tubes. Tubes were vortexed for 2 minutes on medium speed and soil was pelleted at 4C° (2.5KxG for 5 minutes); roots were removed and the soil was pelleted again. Pelleted samples, with supernatant removed, were immediately frozen on dry ice and stored at −80C°. Bulk soil was processed in the same way but without the root removal.

### RNA Extraction

RNA was extracted from 0.5 g of frozen soil using a phenol-chloroform extraction protocol (74) (with modifications from (75)), followed by the Qiagen AllPrep kit to separate DNA from RNA. RNA was treated with TURBO DNase (Thermo Fisher Scientific) following the manufacturer’s protocol to remove residual DNA, and was concentrated using an ethanol precipitation. RNA quality was checked on an Experion Automated Electrophoresis System (Bio-Rad), and quantified using the Qubit RNA BR Assay Kit (Thermo Fisher Scientific).

### Sequencing

Metatranscriptome libraries were prepared and sequenced at the Joint Genome Institute. Ribosomal RNA was depleted from 1 µg of total RNA using the Ribo-Zero rRNA Removal Kit (Epicentre) for Plants and Bacteria. Stranded cDNA libraries were generated using the Illumina TruSeq Stranded RNA LT kit. The rRNA depleted RNA was fragmented and reversed transcribed using random hexamers and SSII (Invitrogen) followed by second strand synthesis. The fragmented cDNA was treated with end-pair, A-tailing, adapter ligation, and 10 cycles of PCR. qPCR was used to determine the concentration of the libraries. The prepared libraries were quantified using KAPA Biosystem’s next-generation sequencing library qPCR kit and run on a Roche LightCycler 480 real-time PCR instrument. The quantified libraries were then multiplexed into pools of 1-3 libraries each, and the pool was then prepared for sequencing on the Illumina HiSeq sequencing platform utilizing a TruSeq paired-end cluster kit, v3, and Illumina’s cBot instrument to generate a clustered flowcell for sequencing. Sequencing of the flowcell was performed on the Illumina HiSeq2000 sequencer using a TruSeq SBS sequencing kit, v3, following a 2×150 indexed run recipe.

### Sequence analysis

Reads were trimmed using Sickle (https://github.com/najoshi/sickle) and BBtools (https://sourceforge.net/projects/bbmap/) was used to remove Illumina adapters and trace contaminants. The reads from all 48 samples were assembled individually using IDBA-UD with default settings (76). Genes were predicted using Prodigal in the anonymous mode (77).

To find the host marker genes, Cox1 and rpS3, we used an HMM profile from Pfam (78), Cox1 (PF00115), and HMMs for Bacteria and Archaea (https://github.com/AJProbst/rpS3_trckr) and searched using hmmsearch (-E 0.00001) from the HMMER suit (79). The identified proteins were classified using both NCBI blast and by making trees. Once classified we predicted genes again using Prodigal in the single mode with the appropriate translation table. The Cox1 protein sequences were dereplicated and clustered at 98% AAI representing an estimated species level designation (80, 81). Assembly errors were found and fixed in the scaffolds containing the Cox1 using ra2.py (https://github.com/christophertbrown/fix_assembly_errors) (82). The bacterial and archaeal rps3 genes was clustered at 99% AAI, representing species level differences (83), using USEARCH (-cluster_fast) (84). The Cox1 and rpS3 trees were generated using references from NCBI, the protein sequences were then aligned using MAFFT v7.402 (85) with the E-INS-i option on Cipres (86). Then alignments were trimmed for the conserved domain manually on Geneious and automatedly trimmed using trimAl (87, 88). The tree was made using RAxML (89) with the JTT protein substitution model (90). The trees were analyzed and figures were generated using iTOL (90). The 18s genes were found and the alignment was generated using ssu_tree.py (https://github.com/christophertbrown/bioscripts27) which searches for rRNA genes using an hmm method then the sequences were dereplicated and clustered at 98% nucleic acid identity, again representing a possible species level designation (91, 92), and aligned using SSU-ALIGN (93). Assembly errors were found and fixed in the scaffolds containing the 18S using ra2.py (https://github.com/christophertbrown/fix_assembly_errors) (82). The tree was generated using the approach described above.

We identified the RNA viral scaffolds using a combined profile HMM approach, using HMMs from Pfam (78) for different types of RdRps. The *Leviviridae* were identified using RNA_replicase_B (PF03431) and scaffolding errors were identified with ra2.py. We used many publicly available HMMs to find the RdRp of eukaryotic RNA viruses. The RdRp genes were initially classified using a blast and tree building method to determine the correct translation table to use for Prodigal in the single mode (77). For mitoviruses we used translation table 4, which to our knowledge is used by all mitoviruses. However, the additional mitoviral genes we predicted may have been incorrectly called if these genomes use a modified genetic code, as occurs in some fungi (94, 95). We examined gene predictions for indications of this (e.g., interruption of the RdRp gene) but as this phenomenon was not identified, no alternative codes were used for gene prediction. RdRp amino acid sequences were dereplicated and clustered at 99% amino acid identity using USEARCH (84). We used previously published alignments for many viral families (23) and added our sequences and key references using the Mafft v7.402 with the --seed (85) and E-INS-i options on Cipres (86). For viral clades without published alignments or where the published alignment was inappropriate (*Fusariviridae, Narnaviridae, Leviviridae* and *Cystoviridae*) we generated our own alignments using reference sequences from NCBI and the same alignment and tree building steps as described above. To identify lysis proteins in the *Leviviridae* genomes we used the Geneious ORF prediction to find all possible open reading frames. The amino acid sequences were run through PSORTb v3.0.2 with the gram negative setting to find possible lysis protein(s) [98].

To obtain coverage values for the viruses we mapped against the entire scaffold containing the RdRp gene. For the presumed hosts we only mapped reads to the open reading frame, for the rpS3 and Cox1. Reads from all samples were mapped using Bowtie2 (--sensitive and --rfg 200,300 options) then reads were filtered for two mismatches using calculate_breadth.py (https://github.com/banfieldlab/mattolm-public-scripts/blob/master/calculate_breadth.py) (97). Coverage values were converted to read counts and normalized with DESeq2 (98). Ordinations were generated from DESeq2 normalized count data in R. The data was ordinated using non-metric multidimensional scaling (R package: vegan) and significantly different clusters were determined using adonis (99). Using DESeq2 we determined the viral and eukaryotic habitat enrichments in the treatments (rhizosphere and litter, bulk and litter and rhizosphere) relative to bulk soil at each time point. We also conducted the opposite test, identifying enrichments in bulk soil by comparing normalized abundance to each treatment (rhizosphere and litter, bulk and litter and rhizosphere) at each time point. P values were corrected for multiple comparisons using a Benjamini-Hochberg correction.

