## Supplemental Figures and Captions for "Metatranscriptomic reconstruction reveals RNA viruses with the potential to shape carbon cycling in soil"

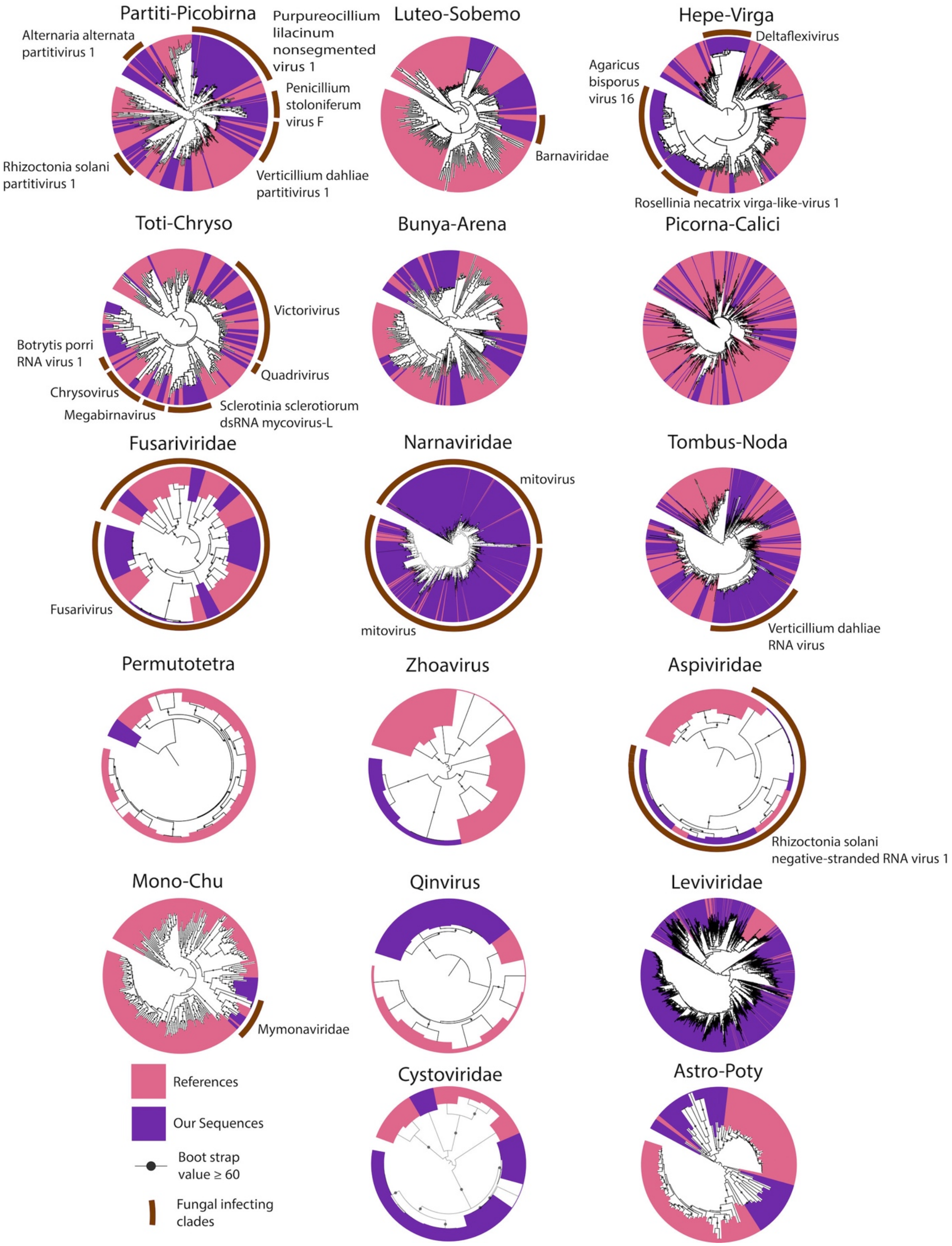


**Supplementary Figure 1.** Trees from Figure 1 with expected fungal infecting clades highlighted in brown. Each predicted fungal infecting clade is labelled with the name of the clade or the name of a well-studied representative virus if the clade is unnamed.

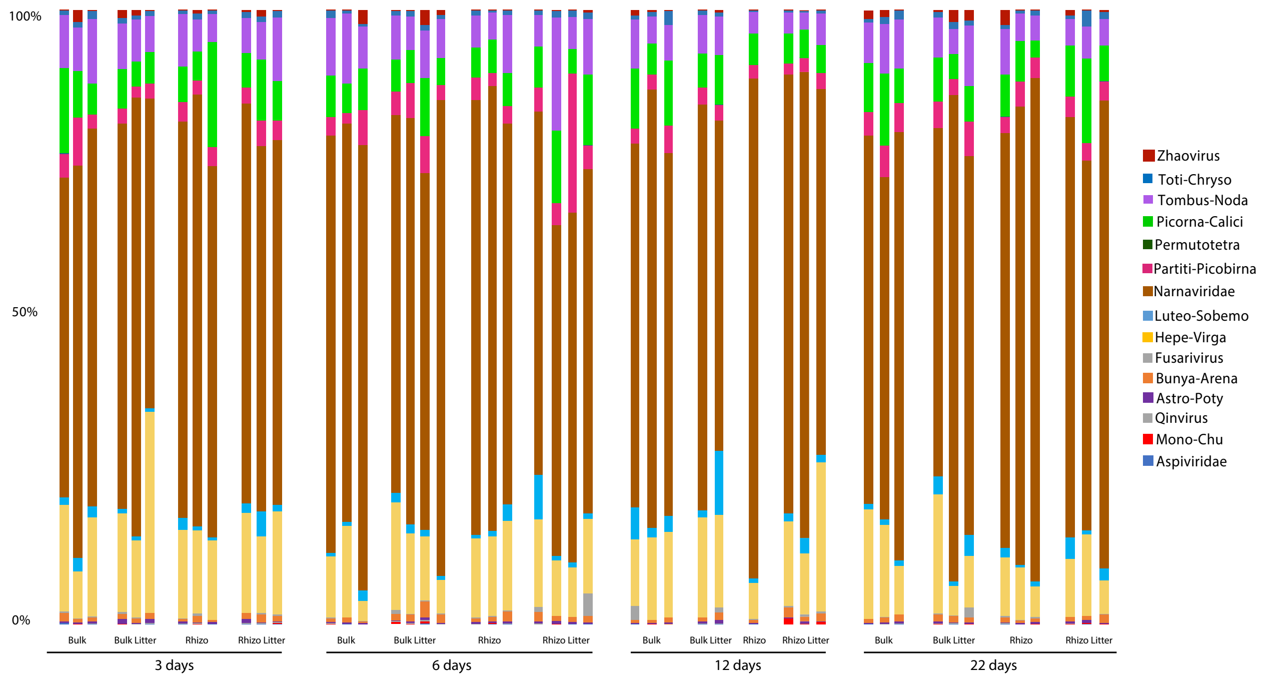


**Supplementary Figure 2.** Percent relative abundance of each eukaryotic RNA viral clade across samples.

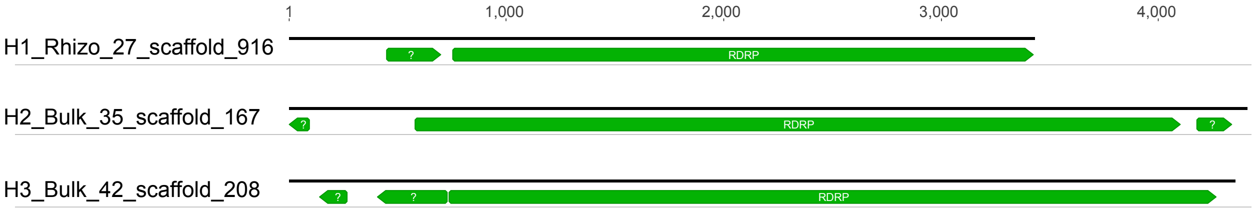


**Supplementary Figure 3.** Representative genome organization identified in some mitoviral genomes.

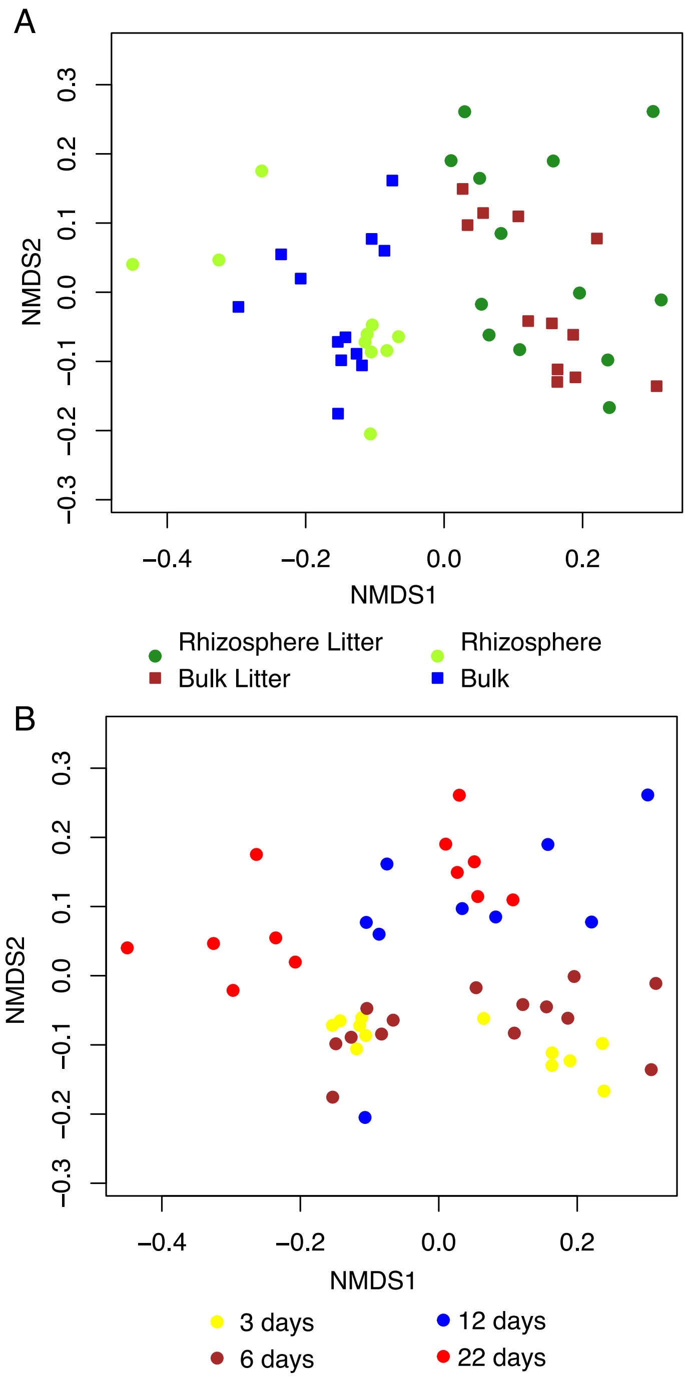


**Supplementary Figure 4.** Nonmetric multidimensional scaling ordination of eukaryotic communities based on Cox1 gene coverage with plant sequences removed. Colored by treatment (A) and harvest day (B).

| Time (d) | Rhizo litter vs Bulk 3 | 6 | 12 | 22 | Bulk litter vs Bulk 3 | 6 | 12 | 22 | Rhizo vs Bulk 3 | 6 | 12 | 22 |
|---|---|---|---|---|---|---|---|---|---|---|---|---|
| Amoebozoa | 43 | 31 | 25 | 36 | 36 | 34 | 21 | 31 | 21 | 21 | 11 | 19 |
| Fungi | 18 | 20 | 19 | 24 | 17 | 24 | 12 | 20 | 15 | 16 | 9 | 18 |
| Unk | 20 | 20 | 12 | 21 | 12 | 28 | 12 | 20 | 14 | 17 | 1 | 5 |
| Metazoa | 16 | 16 | 15 | 13 | 13 | 16 | 10 | 8 | 13 | 13 | 6 | 10 |
| Stramenopiles | 14 | 11 | 10 | 15 | 9 | 12 | 8 | 13 | 3 | 4 | 1 | 2 |
| Alveolata | 10 | 9 | 4 | 3 | 7 | 12 | 4 | 3 | 8 | 9 | 0 | 5 |
| Nucleariidae | 2 | 0 | 0 | 3 | 3 | 1 | 0 | 2 | 0 | 0 | 0 | 1 |
| Heterolobosea | 3 | 3 | 1 | 1 | 1 | 2 | 1 | 1 | 0 | 0 | 1 | 0 |
| Cryptophyta | 3 | 2 | 0 | 2 | 3 | 1 | 0 | 2 | 3 | 0 | 0 | 0 |
| Euglenozoa | 0 | 0 | 0 | 0 | 0 | 0 | 0 | 0 | 0 | 0 | 2 | 1 |
| Viridiplantae | 1 | 1 | 1 | 0 | 0 | 0 | 0 | 0 | 1 | 1 | 1 | 0 |
| Choanoflagellida | 1 | 1 | 0 | 2 | 2 | 1 | 0 | 2 | 2 | 0 | 0 | 0 |
| Alveidia | 2 | 1 | 0 | 0 | 2 | 0 | 1 | 1 | 1 | 0 | 0 | 0 |
| Rhizaria | 0 | 1 | 0 | 1 | 0 | 1 | 0 | 1 | 0 | 1 | 0 | 1 |
| Apusozoa | 0 | 0 | 0 | 0 | 0 | 0 | 0 | 0 | 0 | 0 | 0 | 0 |

| Time (d) | Bulk vs Rhizo litter 3 | 6 | 12 | 22 | Bulk vs Bulk litter 3 | 6 | 12 | 22 | Bulk vs Rhizo 3 | 6 | 12 | 22 |
|---|---|---|---|---|---|---|---|---|---|---|---|---|
| Amoebozoa | 6 | 6 | 6 | 17 | 6 | 8 | 7 | 4 | 9 | 7 | 8 | 16 |
| Fungi | 15 | 14 | 10 | 38 | 7 | 5 | 10 | 8 | 14 | 7 | 9 | 9 |
| Unk | 4 | 4 | 3 | 28 | 2 | 2 | 8 | 6 | 7 | 3 | 5 | 14 |
| Metazoa | 3 | 3 | 5 | 5 | 2 | 4 | 3 | 4 | 7 | 3 | 8 | 3 |
| Stramenopiles | 1 | 7 | 6 | 10 | 3 | 3 | 3 | 0 | 1 | 1 | 0 | 5 |
| Alveolata | 1 | 2 | 3 | 13 | 2 | 1 | 1 | 5 | 7 | 3 | 3 | 10 |
| Nucleariidae | 0 | 0 | 0 | 1 | 0 | 1 | 1 | 0 | 1 | 0 | 0 | 0 |
| Heterolobosea | 0 | 0 | 0 | 2 | 0 | 0 | 0 | 0 | 0 | 0 | 1 | 0 |
| Cryptophyta | 0 | 0 | 0 | 3 | 1 | 0 | 1 | 1 | 1 | 1 | 0 | 1 |
| Euglenozoa | 0 | 0 | 0 | 1 | 0 | 0 | 0 | 0 | 0 | 0 | 0 | 0 |
| Viridiplantae | 1 | 1 | 0 | 0 | 0 | 2 | 0 | 1 | 0 | 0 | 1 | 0 |
| Choanoflagellida | 0 | 0 | 0 | 2 | 0 | 0 | 0 | 0 | 0 | 0 | 0 | 1 |
| Alveidia | 0 | 0 | 0 | 0 | 0 | 0 | 0 | 0 | 0 | 0 | 0 | 0 |
| Rhizaria | 0 | 0 | 1 | 0 | 0 | 0 | 0 | 1 | 0 | 0 | 0 | 0 |
| Apusozoa | 0 | 0 | 0 | 0 | 0 | 0 | 0 | 0 | 0 | 0 | 0 | 0 |

**Supplementary Figure 5.** Time-series heatmap representing the enriched clades of eukaryotes. The enrichment was determined by comparing the Cox1 transcript level of the treatments against the bulk soil (A) and the bulk soil against the treatments (B). Red indicates a higher number of identified significant enrichments and blue indicates no species were significantly enriched. The clades of eukaryotes are ordered by the total number of species identified.

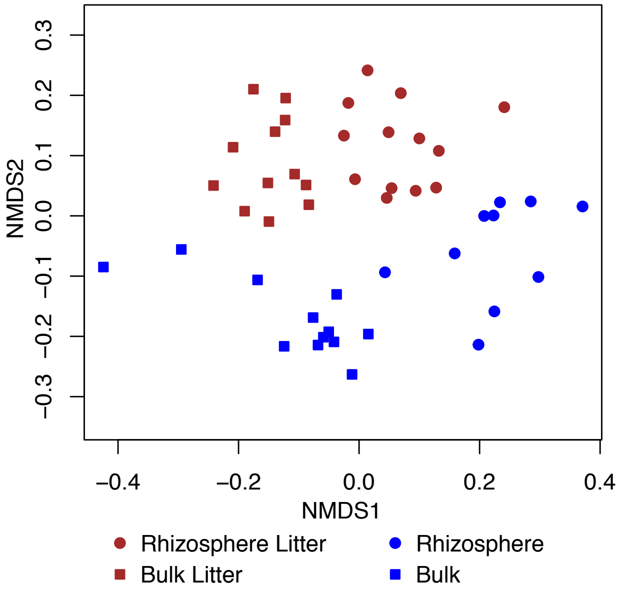


**Supplementary Figure 6.** Nonmetric multidimensional scaling ordination of eukaryotic RNA viral community with *Narnaviridae* sequences removed.

| Time (d) | Rhizo litter vs Bulk 3 | 6 | 12 | 22 | Bulk litter vs Bulk 3 | 6 | 12 | 22 | Rhizo vs Bulk 3 | 6 | 12 | 22 |
|---|---|---|---|---|---|---|---|---|---|---|---|---|
| Narnaviridae | 2 | 1 | 9 | 25 | 2 | 1 | 2 | 10 | 0 | 0 | 24 | 13 |
| Tombus-Noda | 2 | 3 | 2 | 2 | 0 | 1 | 0 | 0 | 0 | 0 | 0 | 1 |
| Parti-picobirna | 0 | 2 | 0 | 0 | 0 | 0 | 0 | 0 | 0 | 0 | 0 | 0 |
| Hepe-Virga | 0 | 1 | 1 | 0 | 0 | 1 | 1 | 0 | 0 | 0 | 0 | 0 |
| Picorna-Calici | 0 | 2 | 2 | 4 | 1 | 1 | 0 | 1 | 0 | 1 | 0 | 1 |
| Bunya-Arena | 2 | 0 | 1 | 1 | 1 | 1 | 1 | 1 | 0 | 0 | 0 | 0 |
| Toti-Chryso | 1 | 1 | 0 | 3 | 0 | 0 | 0 | 1 | 0 | 0 | 0 | 0 |
| Luteo-Sobemo | 0 | 0 | 0 | 3 | 0 | 0 | 4 | 2 | 0 | 0 | 0 | 0 |
| Astro-Poty | 0 | 2 | 2 | 2 | 0 | 0 | 0 | 0 | 0 | 0 | 0 | 0 |
| Mono-Chu | 0 | 0 | 0 | 0 | 0 | 0 | 0 | 0 | 0 | 0 | 0 | 0 |
| Fusarivirus | 0 | 0 | 0 | 0 | 0 | 0 | 0 | 0 | 0 | 0 | 0 | 0 |
| Aspiviridae | 0 | 0 | 0 | 0 | 0 | 0 | 0 | 0 | 0 | 0 | 0 | 0 |
| Qinvirus | 0 | 0 | 0 | 0 | 0 | 0 | 0 | 0 | 0 | 0 | 0 | 0 |
| Zhaovirus | 0 | 0 | 0 | 0 | 0 | 0 | 0 | 0 | 0 | 0 | 0 | 0 |
| Permutotetra | 0 | 0 | 0 | 0 | 0 | 0 | 0 | 0 | 0 | 0 | 0 | 0 |

| Time (d) | Bulk vs Rhizo litter 3 | 6 | 12 | 22 | Bulk vs Bulk litter 3 | 6 | 12 | 22 | Bulk vs Rhizo 3 | 6 | 12 | 22 |
|---|---|---|---|---|---|---|---|---|---|---|---|---|
| Narnaviridae | 4 | 21 | 0 | 3 | 1 | 9 | 0 | 0 | 0 | 0 | 1 | 0 |
| Tombus-Noda | 5 | 2 | 0 | 4 | 3 | 0 | 0 | 1 | 1 | 0 | 2 | 1 |
| Parti-picobirna | 0 | 0 | 0 | 0 | 0 | 0 | 0 | 0 | 0 | 0 | 5 | 0 |
| Hepe-Virga | 1 | 0 | 1 | 0 | 0 | 0 | 0 | 0 | 0 | 0 | 2 | 1 |
| Picorna-Calici | 0 | 0 | 0 | 2 | 0 | 0 | 1 | 1 | 0 | 0 | 1 | 2 |
| Bunya-Arena | 0 | 0 | 0 | 1 | 0 | 0 | 0 | 1 | 0 | 0 | 0 | 0 |
| Toti-Chryso | 0 | 0 | 0 | 1 | 0 | 0 | 0 | 0 | 0 | 0 | 7 | 0 |
| Luteo-Sobemo | 0 | 0 | 0 | 0 | 0 | 0 | 0 | 0 | 0 | 0 | 1 | 0 |
| Astro-Poty | 0 | 0 | 0 | 0 | 0 | 0 | 0 | 0 | 0 | 0 | 0 | 0 |
| Mono-Chu | 0 | 0 | 0 | 0 | 0 | 0 | 0 | 0 | 0 | 0 | 0 | 0 |
| Fusarivirus | 0 | 0 | 0 | 0 | 0 | 0 | 0 | 0 | 0 | 0 | 0 | 0 |
| Aspiviridae | 0 | 0 | 0 | 0 | 0 | 0 | 0 | 0 | 0 | 0 | 0 | 0 |
| Qinvirus | 0 | 0 | 0 | 0 | 0 | 0 | 0 | 0 | 0 | 0 | 2 | 0 |
| Zhaovirus | 0 | 0 | 1 | 1 | 0 | 0 | 1 | 1 | 0 | 0 | 1 | 1 |
| Permutotetra | 0 | 0 | 0 | 0 | 0 | 0 | 0 | 0 | 0 | 0 | 0 | 0 |

**Supplementary Figure 7.** Time-series heatmap representing the enriched clades of eukaryote viruses. The enrichment was determined by comparing the viral transcript level of from the treatments against the bulk soil (A) and the bulk soil against the treatments (B). Red indicates a higher number of identified significant enrichments and blue indicates no species were significantly enriched. The clades of viruses are ordered by the total number of sequences identified.

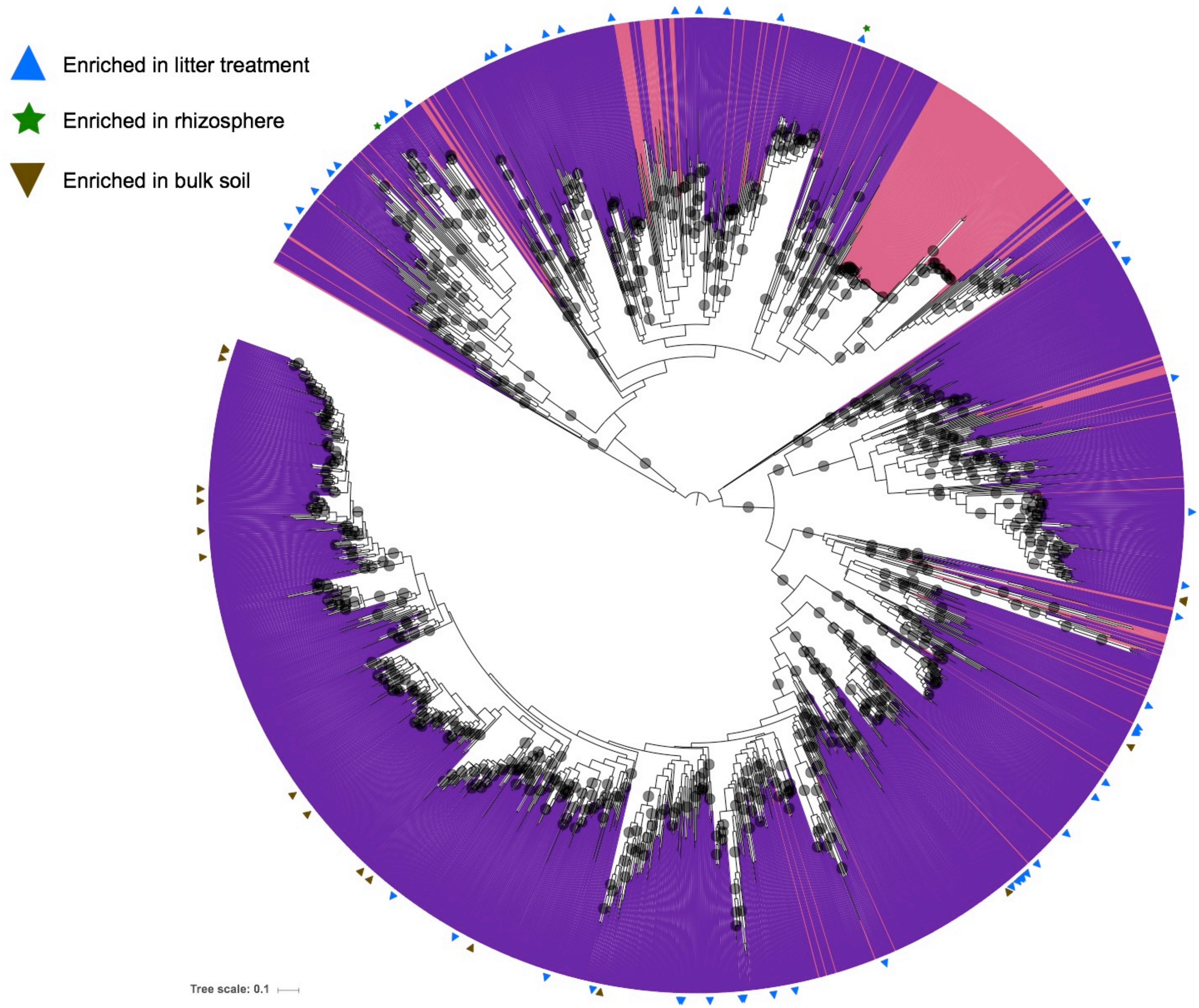


**Supplementary Figure 8.** Statistically enriched *Leviviridae* in the treatments. Blue triangles represent *Leviviridae* that were enriched in the treatments samples (rhizosphere and litter, bulk and litter) compared to unamended bulk soil. Green stars mark viral strains that were enriched in the unamended rhizosphere compared to bulk soil. And brown triangles represent *Leviviridae* enriched in bulk soil when compared to the treatment samples.

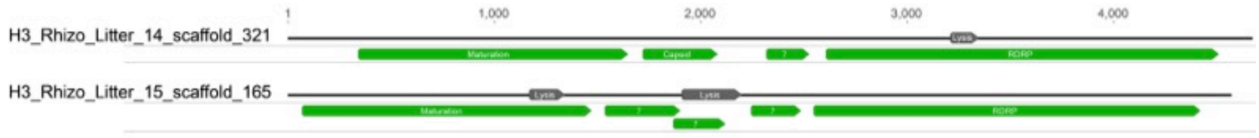


**Supplementary Figure 9.** Representative novel genome architectures identified in *Leviviridae.*

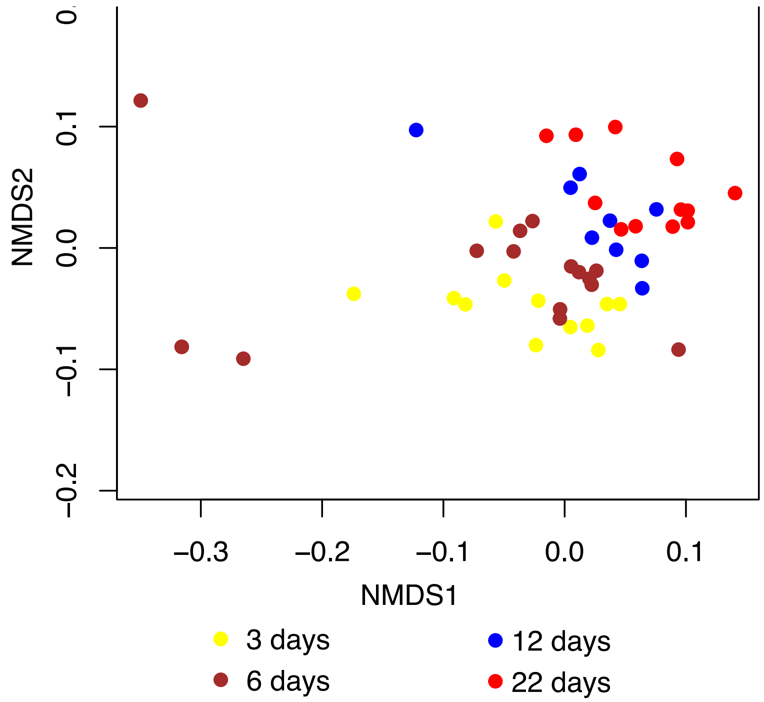


**Supplementary Figure 10.** Nonmetric multidimensional scaling ordination of *Leviviridae* community colored by harvest day.

**Supplementary Data 1.** Tree files for each viral family.
**Supplementary Data 2.** Eukaryotic COX1 tree files.
**Supplementary Data 3.** 18S rRNA tree files.
**Supplementary Data 4.** Proteobacteria rpS3 tree files
